# The essential role of hypermutation in rapid adaptation to antibiotic stress

**DOI:** 10.1101/422642

**Authors:** Heer H. Mehta, Amy G. Prater, Kathryn Beabout, Ryan A. L. Elworth, Mark Karavis, Henry S. Gibbons, Yousif Shamoo

## Abstract

A common outcome of antibiotic exposure in patients and *in vitro* is the evolution of a hypermutator phenotype that enables rapid adaptation by pathogens. While hypermutation is a robust mechanism for rapid adaptation, it requires trade-offs between the adaptive mutations and the more common “hitchhiker” mutations that accumulate from the increased mutation rate. Using quantitative experimental evolution, we examined the role of hypermutation in driving adaptation of *Pseudomonas aeruginosa* to colistin. Metagenomic deep sequencing revealed 2,657 mutations at ≥ 5% frequency in 1,197 genes and 761 mutations in 29 end point isolates. By combining genomic information, phylogenetic analyses, and statistical tests, we showed that evolutionary trajectories leading to resistance could be reliably discerned. In addition to known alleles such as *pmrB,* hypermutation allowed identification of additional adaptive alleles with epistatic relationships. Although hypermutation provided a short-term fitness benefit, it was detrimental to overall fitness. Alarmingly, a small fraction of the colistin adapted population remained colistin susceptible and escaped hypermutation. In a clinical population, such cells could play a role in re-establishing infection upon withdrawal of colistin. We present here a framework for evaluating the complex evolutionary trajectories of hypermutators that applies to both current and emerging pathogen populations.

**Importance**

Bacteria can increase mutation rates in response to stress as an evolutionary strategy to avoid extinction. However, the complex mutational landscape of hypermutators makes it difficult to distinguish truly adaptive mutations from hitchhikers that follow similar evolutionary trajectories. We provide a framework for evaluating the complex evolutionary trajectories of hypermutators that can be applied to both current and emerging pathogen populations. Using *Pseudomonas aeruginosa* evolving to colistin as a model system, we examine the essential role of hypermutation in the evolution of resistance. Additionally, our results highlight the presence of a subset of cells that survive and remain susceptible during colistin exposure which can serve as a reservoir for re-infection upon withdrawal of the drug in clinical infections. This study provides a broad understanding of hypermutation during adaptation and describes a series of analyses that will be useful in identifying adaptive mutations in well annotated and novel bacterial mutator populations.

## Introduction

Hypermutation is a phenomenon that is often observed in clinical isolates of pathogenic species (1–3). The opportunistic pathogen, *Pseudomonas aeruginosa* is a common nosocomial pathogen affecting immunocompromised patients, especially those with cystic fibrosis (CF). 30 to 54% of CF patients infected with *P. aeruginosa* are colonized by hypermutator strains that are associated with reduced lung function and chronic infections (4–6). During infection, *P. aeruginosa* also forms persistent and difficult to clear biofilms that are known to exhibit reduced susceptibility to antimicrobials (7, 8). *P. aeruginosa* also exhibits great metabolic and genetic plasticity allowing it to readily acquire resistance to antibiotics (9). Resistance to the drug of last resort, a cationic antimicrobial peptide (CAP) called colistin (Polymyxin E), has been observed and has been associated with hypermutation (10–13).

Mutation acquisition during selection is dependent on mutation supply which can be boosted by hypermutation (14, 15). The typical path leading to a hypermutator phenotype is mutations in the DNA repair system including the MutS/MutL class of proteins (14, 16, 17). The increase in mutation supply accelerates the rate at which bacteria become resistant and thus, hypermutators explore the adaptive evolutionary trajectories leading to resistance more rapidly than non-hypermutators. Interestingly, the dramatic increase in mutation rate also means that many non-adaptive mutations accumulate and are carried along as “hitchhikers” that in the long run are likely to decrease the overall fitness of the organisms in non-selective conditions (18, 19). Nevertheless, hypermutation provides a swift strategy for cells undergoing stress to acquire adaptive mutations before they go extinct.

Experimental evolution is a powerful approach to understanding the genetic and biochemical basis for antibiotic resistance (16, 20–25). In this work, *P. aeruginosa* PAO1 was evolved to colistin as a continuous culture in a bioreactor where the population was constantly maintained at mid-exponential phase in the presence of sub-inhibitory drug concentrations (24). The observation of hypermutation, accompanied by a dramatic increase in mutation rate among the evolving PAO1 population presented both challenges and opportunities for understanding the evolution of colistin resistance. The potential benefits of this complex data set, however, were equally clear. Hypermutation generated an extensive, if not nearly exhaustive survey of the accessible evolutionary trajectories leading to colistin resistance.

We used a combination of phylogenetic and statistical approaches to sort through these complex data, discern truly adaptive alleles from hitchhikers and infer not just the major genetic changes associated with colistin resistance but also those alleles that, in combination with the major players were essential to produce the high minimum inhibitory concentrations (MICs) observed in many clinical isolates (12, 26). Our work has benefited strongly from a series of clinical and *in vitro* experimental evolution studies that have identified many of the major contributors to colistin resistance in *P. aeruginosa* (10, 12, 13, 26, 27).

Taken together, our work provides a comprehensive survey of the alleles responsible for *P. aeruginosa* resistance to colistin and perhaps, more importantly, a means to examine future hypermutators in less well characterized or emerging pathogens. The increased mutation rate of hypermutators provides a major adaptive advantage to this pathogen during exposure to colistin. This highlights the strength of hypermutation as an adaptive strategy during exposure to stress and the adaptability of *P. aeruginosa* to survive under such conditions, making it a formidable pathogen.

## Results

### Hypermutator variants of *P. aeruginosa* emerge rapidly after colistin exposure during continuous experimental evolution

*P. aeruginosa* PAO1 was evolved to colistin as a continuous culture in a bioreactor using quantitative experimental evolution (24). The starting MIC of colistin for PAO1 was 1-2 mg/l. Over the course of experiment, the PAO1 population was exposed to increasing, but sub-inhibitory, concentrations of colistin to a final concentration of 16 mg/l colistin, which is four times higher than the clinical breakpoint for resistance of *P. aeruginosa* to colistin (MIC ≥ 4mg/l) (28). Genomic DNA from each daily population was prepared for metagenomic deep sequencing. At the end of the experiment, the final population was serially diluted and spread on a non-selective growth medium for the isolation of end point isolates for whole genome sequencing. The end point isolates produced colonies of diverse morphologies (Supplementary file S1) consistent with the selection of a highly polymorphic population within the bioreactor experiments and this was consistent across duplicate experiments. Single colonies isolated from the final populations ranged from being totally susceptible to totally resistant to colistin which is also consistent with a diverse polymorphic population (Fig. S1 (b) and Table S1). 29 end point isolates were selected for whole genome sequencing.

Of the 29 sequenced isolates, 25 contained mutations within *mutS,* the gene encoding the DNA mismatch repair enzyme MutS (Fig. 1).Those end point isolates that had acquired *mutS* mutations had from 44 to 92 mutations each while strains without mutations in *mutS* had 5 to 9 mutations showing that mutations within *mutS* correlated with increased genetic diversity. Metagenomic deep sequencing of daily populations showed that on day 10 of adaptation during both, runs 1 and 2 (corresponding to 1.75 mg/l colistin in run 1 and 2 mg/l colistin in run 2), mutations were observed readily in *mutS* (Fig. 1). While the majority of the population contained mutations within *mutS*, a fraction of the final population did not become hypermutators (14.6% in run 1 and 7.6% in run2).

**Figure 1.**
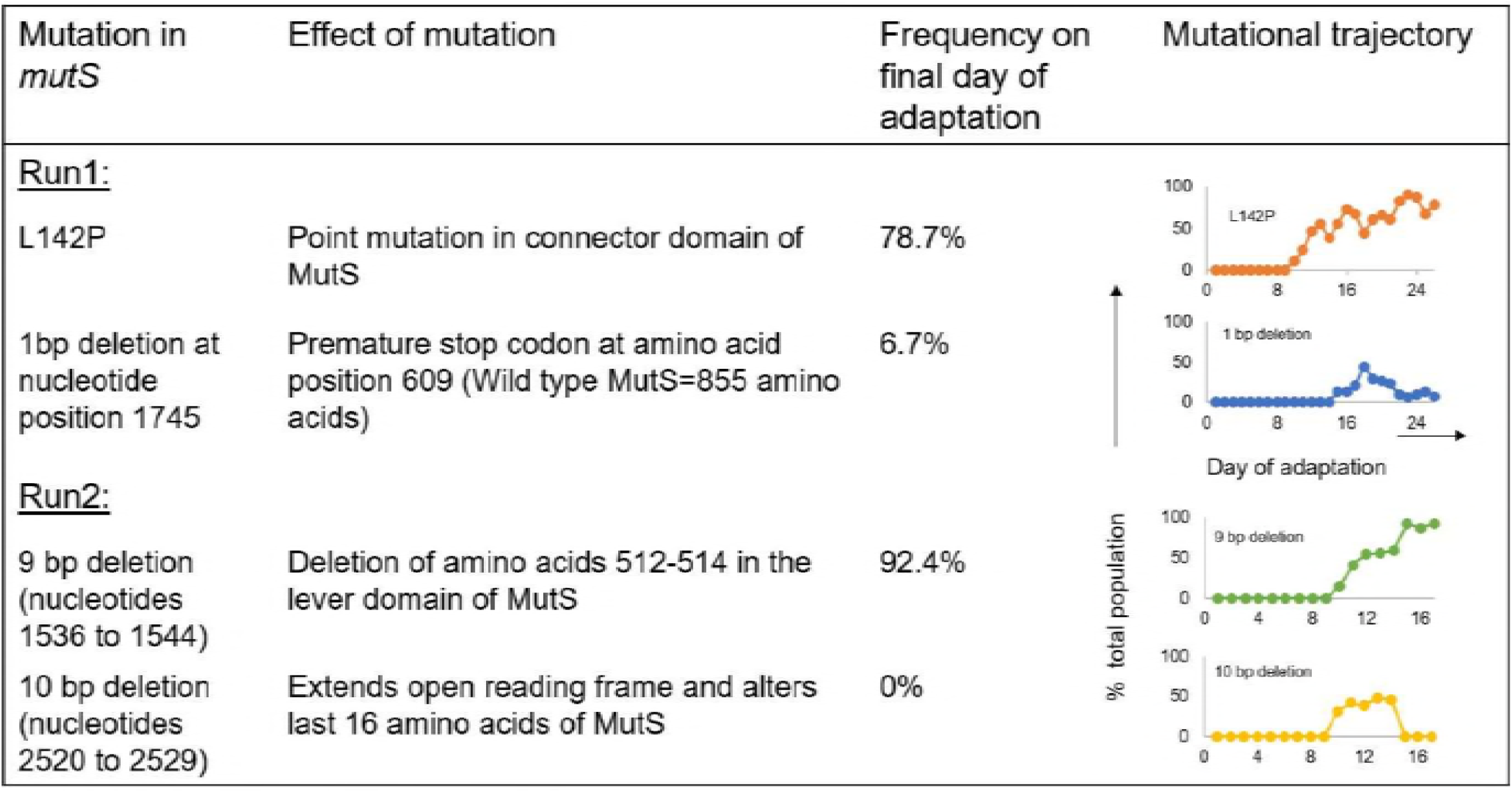
*mutS* mutations observed during adaptation of *P. aeruginosa* populations to colistin. In duplicate experiments (Run 1 and Run 2), mutations were seen in the *mutS* gene. The specific mutations identified, their effect on MutS and their abundance in the evolved populations are indicated (mutation detection cut-off = 5%). The last column shows the frequency for each *mutS* mutation over the course of evolution. The X-axis shows the day of adaptation (26 days in Run 1 and 17 in Run 2) and the Y-axis indicates the percentage of the total population that possessed the mutation.

As expected, the acquisition of the *mutS* mutations was accompanied by a rapid increase in mutation frequency in the total evolving population. As seen in Figure 2, the starting population on day 1 of evolution, in both runs, had a basal level of diversity. Aside from the underlying diversity in the population, what is apparent from Figure 2 is that as soon as *mutS* mutations arose in both the evolving populations (indicated by red stars), the populations started accumulating more mutations that rose to higher frequencies than mutations contributing to the basal diversity in the population. This was accompanied by increased non-susceptibility of the total population to colistin. Interestingly, lineages derived from Run 1 *mutS* with a 1 bp deletion that introduces a premature stop codon at residue 609 and Run 2 *mutS* with a 10 bp deletion that alters the last 16 amino acids both go to extinction, demonstrating that hypermutation, while increasing the probability of finding successful evolutionary trajectories, does not guarantee success. It may also be that the unsuccessful *mutS* mutant lineages may have stronger mutator phenotypes that accumulate deleterious mutations more rapidly leading to a faster decrease in overall fitness compared to the more successful *mutS* mutants in Runs 1 and 2.

**Figure 2.**
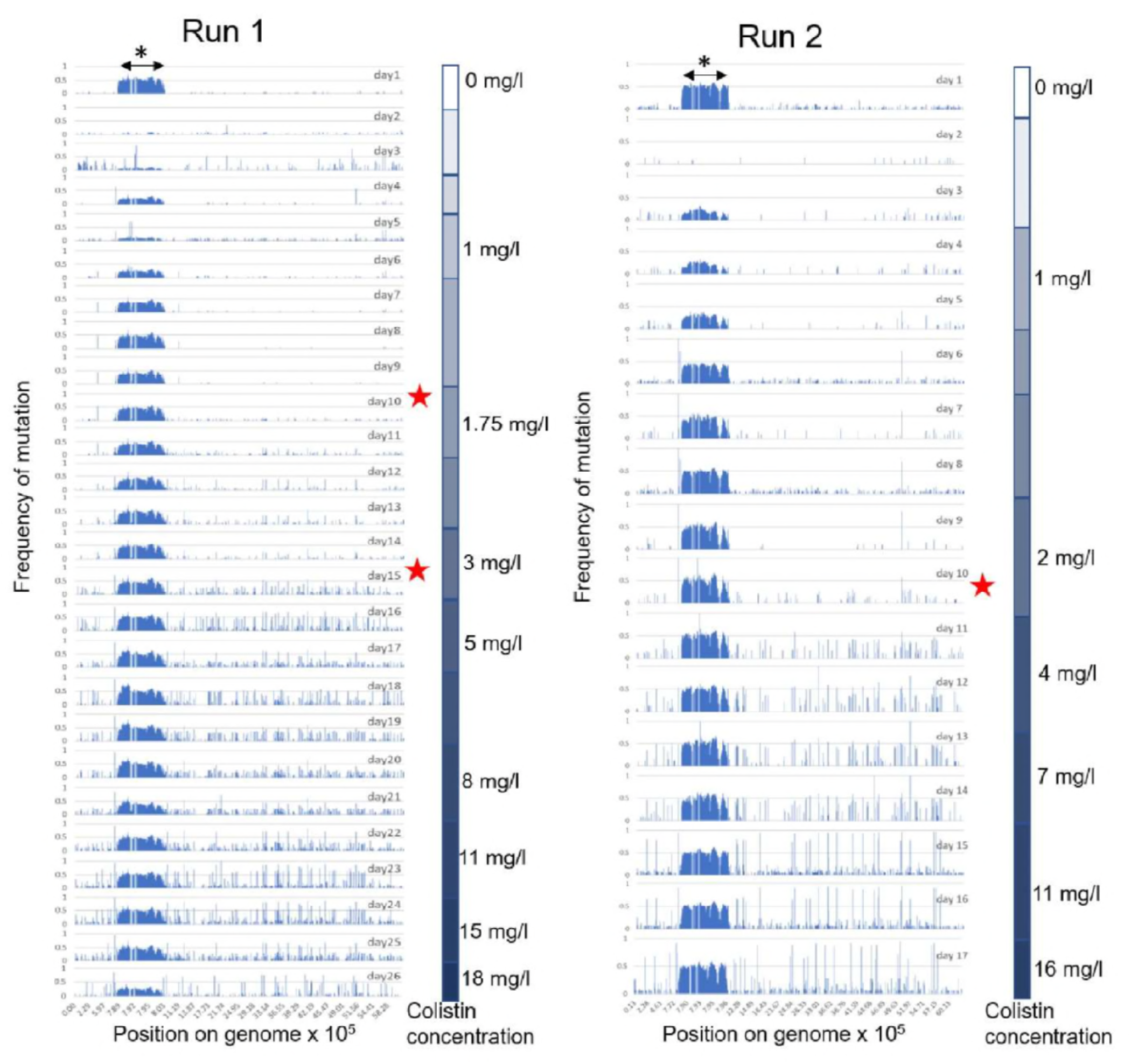
Single nucleotide polymorphism (SNP) map of the PAO1 populations evolving to colistin (Run 1 and Run 2). For ease of comparison, the graphs represent a linearized genome of PAO1 with SNP positions indicated on the X-axis. The X-axis is not distributed evenly but denotes positions on the genome carrying mutations. This is done to highlight regions of the genome with higher mutation density. The Y-axis is the frequency of the mutation in the total population. For each run of adaptation, graphs of daily sampled populations are stacked (26 days of evolution for Run 1 and 17 days for Run 2). Red stars indicate days on which *mutS* mutations arose in each population. The color gradient on the right side of each panel shows the step-wise increase in colistin concentration experienced by the populations during adaptation. The increased genetic diversity in the region from 7.8×10^5^ to 8×10^5^ bp on the PAO1 genome can be attributed to the Pf4 phage island (indicated by asterisk), details of which are provided in Supplementary file S1.

### Evolutionary relationship of end point isolates highlights the role of secondary mutations in resistance

Phylogenetic trees were constructed to identify the linkages of mutations within the end point isolates (Fig. 3). The 29 sequenced end point isolates had cumulatively acquired 761 total mutations affecting 563 genes. It was interesting to note that in both adaptation runs, isolates that did not acquire *mutS* mutations were phylogenetically closely related to the ancestor and had no increase in colistin resistance. The lack of success in achieving high levels of colistin tolerance by the non-hypermutators highlights the higher efficiency of sampling the potential evolutionary trajectories by strains with elevated mutation rates. End point isolates containing *mutS* mutations had undergone considerable divergence, with each branch varying substantially in total number and type of mutations. It was evident that being a hypermutator alone was not sufficient to acquire resistance. Isolates like I1-6 and I1-76 that had MutS^L142P^ were still susceptible (MIC = 2 mg/l) and thus while mutations to *mutS* are drivers for adaptation they are not directly responsible for increased colistin resistance.

**Figure 3.**
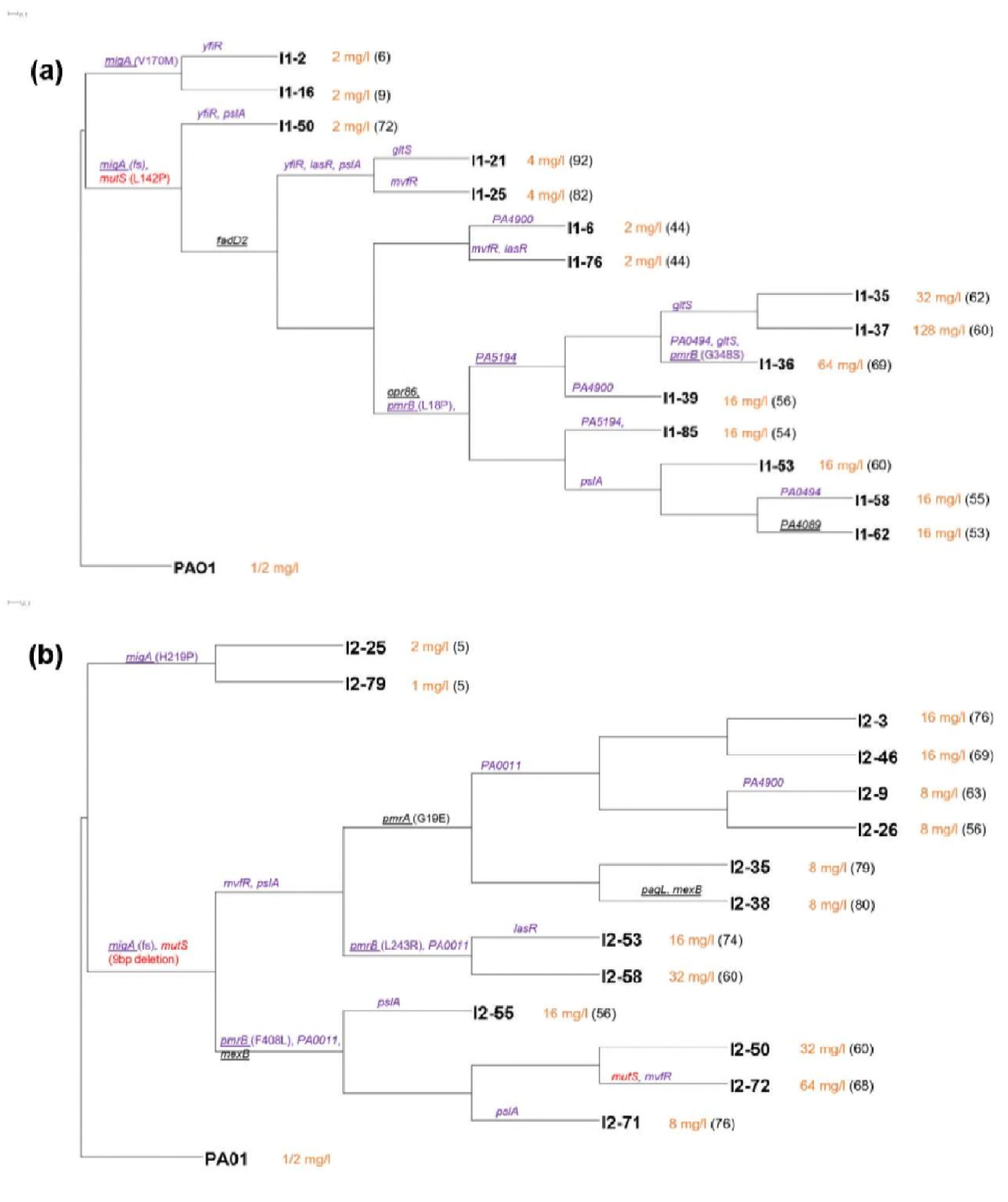
Phylogenetic trees for end point isolates obtained from experimental evolution Runs 1 (a) and 2 (b). PAO1: Ancestor. Isolate names are at the right side of each branch. Orange text following the isolate name denotes its colistin MIC and in parenthesis are the total number of mutations identified in that particular isolate. Branches with mutations in putative targets identified in this study have names of mutated genes on them. Targets in purple text were identified by the Fisher’s Exact test of end point isolates and targets underlined were common among our study and other polymyxin resistance studies. *mutS* mutations are identified in red text. The large number of mutations per hypermutator lineage precludes their complete inclusion in these phylogenetic trees. The complete list of mutations in each end point isolate can be found in Datasets S3 and S4.

The highest levels of resistance were achieved by end point isolates that emerged from branches containing mutations in the *pmrAB* genes. Previous studies in *P. aeruginosa* have identified the role of PmrAB in resistance to cationic antimicrobial peptides (CAPs) (29, 30). In Figure 3(a), the branch with the initial *pmrB* mutation diverged into several branches leading to end point isolates that varied in MICs from 16 to 128 mg/l. Similarly, in Figure 3(b), different end point isolates diverging from the same *pmrA* or *pmrB* branch had different colistin MICs. This suggested that although mutations in *pmrAB* were required for achieving resistance, additional mutations in resistant end point isolates were playing an essential role in increasing resistance to colistin.

### Hypermutation reveals challenges in distinguishing adaptive mutations from hitchhikers

As the final populations of experimental evolution were dominated by cells with mutator phenotypes, classical genetic approaches for the validation of adaptive alleles such as reintroducing the proposed changes into a clean genomic background were effectively impossible (with an average of 60 mutations per hypermutator end point isolate, there were 60! possible combinations of mutations) and required a different approach. While hypermutation introduced a large number of non-adaptive hitchhiker alleles into both the metagenomic and end point isolate genomic data, the extensive mutational saturation offered a methodological path forward. A statistical approach was used to identify potentially adaptive genes based on the concept that if a larger than expected number of mutations in the same gene across various end point isolates from both the runs were identified then those genes were more likely to be adaptive (16). Under the null hypothesis that all mutations were randomly distributed across the genome, 11 genes were identified that were mutated more frequently than expected in the end point isolates using the Fisher’s Exact Test (Table 1). A total of 563 genes were mutated in the 29 sequenced end point isolates. By plotting the number of mutations per gene versus the percentage of end point isolates having a mutation in that gene (Fig. 4), it was observed that some of the statistically significant genes (*p* value < 0.001) were located on the top right quadrant. The exception to this was *mutS.* Although *mutS* was seen in 25 of the 29 sequenced end point isolates that have a total of 3 unique mutations in this gene, the *mutS* gene itself was sufficiently long (2568 bp) so that it did not meet the *p* value cut-off of 0.001 for being called significant. If a gene is very long or if a single mutation or small subset of mutations are the only possible adaptive changes in that gene then it may not rise above the required *p* value according to the Fisher’s Exact Test. This highlights a shortcoming of this particular approach and explains why we combine it with other methods (discussed later) to increase the overall success in identifying adaptive alleles.

**Table 1.** Putative targets involved in colistin resistance in PAO1 based on mutations observed in end point isolates

| Gene name | Mutation(s) identified in end point isolates | p-value from Fisher's Exact Test | Function |
| --- | --- | --- | --- |
| **Fisher's Exact Test of end point isolates** |  |  |  |
| *migA* | frameshifts, V170M, H219P | 5.58e-6 | Alpha-1,6-rhamnosyl transferase, responsible for uncapped core oligosaccharides in the LPS (49) |
| *mvfR* | Y77H, V162A, L6P, V9V | 8.39e-6 | Quorum sensing and virulence regulator |
| *pmrB* | L18P, G348S, L243R, F408L | 3.4e-5 | Two component system involved in cationic antimicrobial sensing and resistance (30) |
| *pslA* | frameshifts, 700 bp deletion, G364D, R229C, Y268H | 3.43e-5 | Exopolysaccharide involved in biofilm formation; involved in biofilm resistance to antibiotics (50) |
| *yfiR* | L183P, C71S, Y58C | 5.43e-5 | Part of c-di-GMP regulator system involved in biofilm formation. |
| *lasR* | L118P, G235D, T115A | 1.06e-4 | Quorum sensing regulator |
| *PA5194* | W64*, G227D, W239* | 1.46e-4 | LpxT; phosphorylation of lipid A |
| *PA0011* | frameshifts, W33* | 1.96e-4 | 2-hydroxylauroyl transferase; transfers 2-hydroxylaurate to C-2 position of lipid A |
| *gltS* | A194V, G351S, L126L | 4.86e-4 | Glutamate/sodium symporter |
| *PA4900* | L321P, P339P, Q4* | 6.45e-4 | Probable major facilitator superfamily transporter |
| *PA0494* | A44V, P56L, Y201Y | 6.96e-4 | Probable acyl-CoA carboxylase subunit |
| **Observed in other polymyxin resistance studies of *P. aeruginosa*** |  |  |  |
| *pmrAB* | G19E, L18P, G348S, L243R, F408L |  | Two component system involved in cationic antimicrobial sensing and resistance (12, 13, 27, 30) |
| *migA* | frameshifts, V170M, H219P |  | Alpha-1,6-rhamnosyl transferase, responsible for uncapped core oligosaccharides in the LPS (12, 13) |
| *PA5194* | W64*, G227D, W239* |  | Lipid a kinase; adds phosphate group to lipid A (12) |
| *fadD2* | Q245R |  | Long chain fatty acid CoA ligase (12) |
| *opr86* | D535N |  | Outer membrane protein (12) |
| *PA4089* | G105D |  | Probable short chain dehydrogenase involved in fatty acid biosynthesis (27) |
| *pagL* | D118G |  | Lipid A deacylation, upregulated by polymyxin (34) |
| *mexB* | T242A, T295A |  | RND multidrug efflux transporter (12) |

**Figure 4.**
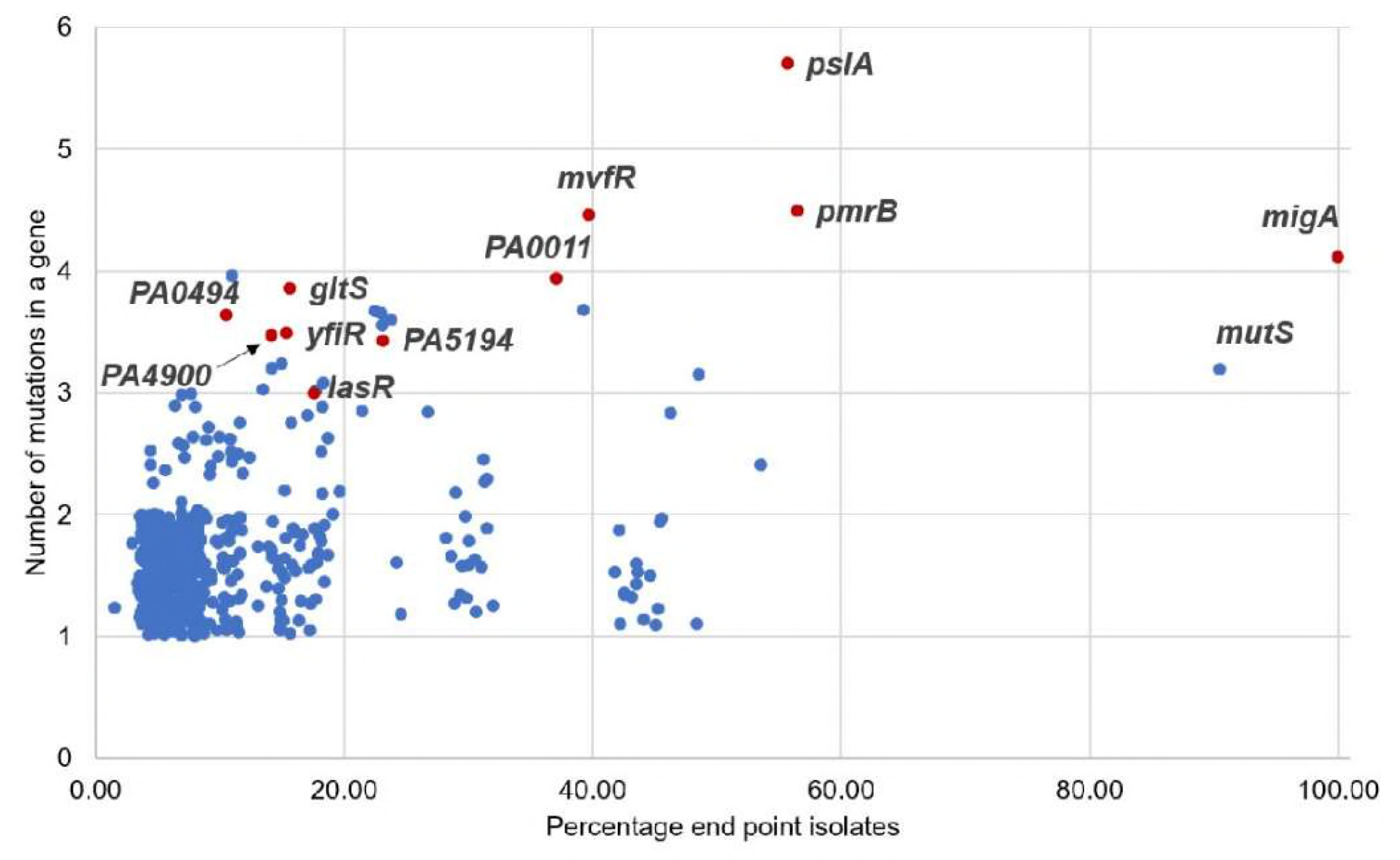
Genes mutated more frequently than expected identified using Fisher’s Exact Test. Number of mutations identified within a single gene among the 29 sequenced end point isolates was plotted against the percentage of end point isolates containing a mutation in that gene. Genes identified as significant with a *p* value less than 0.001 in the Fisher’s Exact Test are highlighted in red. False noise has been added to the data points to separate them on the graph. Detailed analysis of this graph is provided in Supplementary file S1.

Among the evolving daily populations from both runs, 1,197 genes were mutated and a total of 2657 mutations were identified at ≥ 5% frequency. When this same test was performed on these mutations, 41 genes were identified as significant (Table 2). Among these 41 genes, four were also identified in the Fisher’s Exact Test of the end point isolates (*pmrB, PA0011, migA* and *pslA*). *pmrA,* which is known to be involved in resistance, was not identified as a significant gene in the end point isolates but was in the daily populations. The power of this test could certainly be increased by having data from multiple evolving populations instead of the 2 runs conducted in this work. Taken together however, the cumulative data from the 2 populations and 29 end point isolates provides an extensive list of genes that potentially play a role in colistin resistance.

**Table 2:**
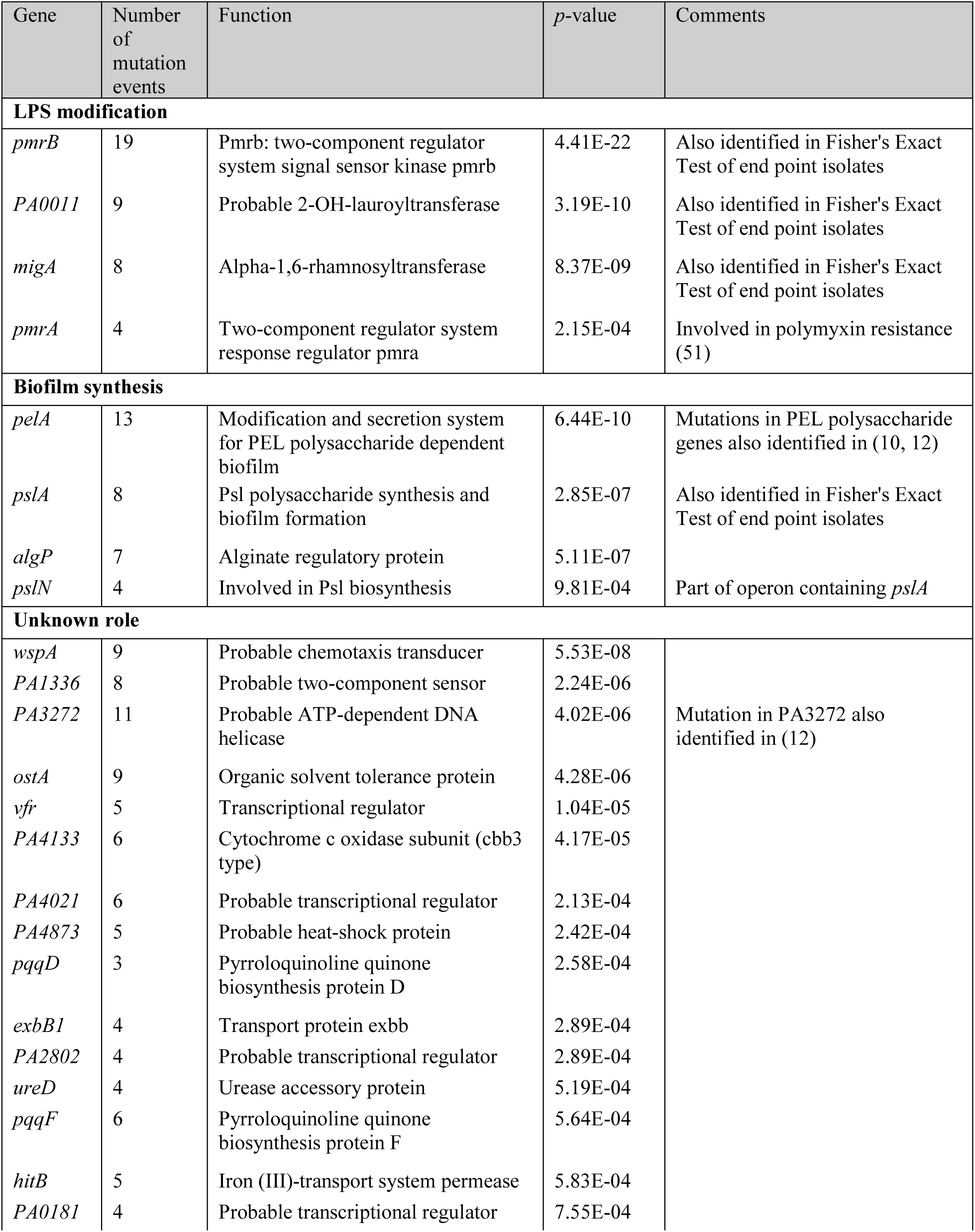
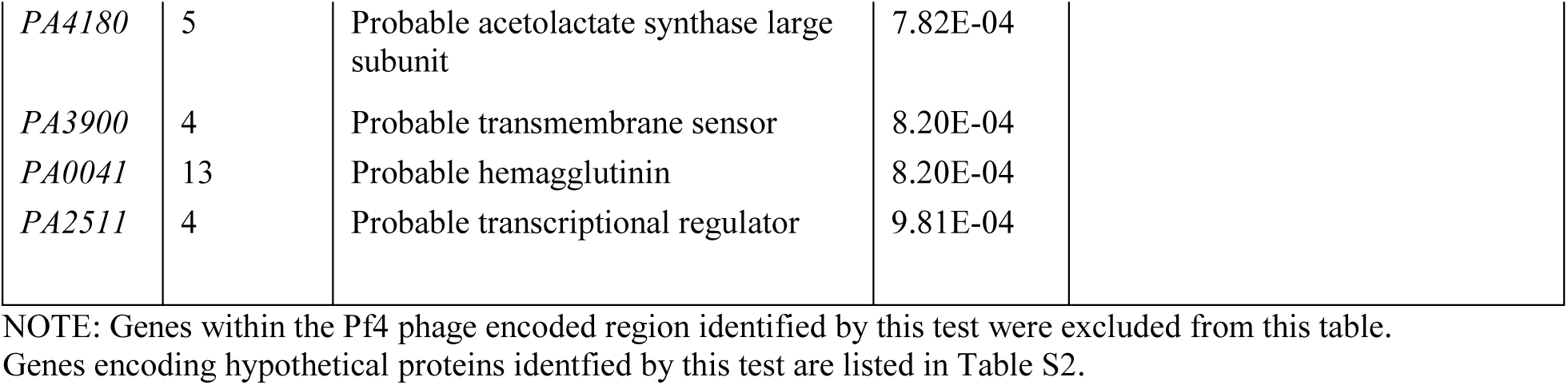
Candidate colistin resistance genes identified as significant by Fisher’s Exact Test of daily populations

### Identification of additional genes associated with colistin resistance

Previous studies have been conducted to identify genetic changes leading to polymyxin resistance in *P. aeruginosa* (11–13, 26, 27, 30–32). Since resistance has been often associated with hypermutation (12, 13) which leads to accumulation of a large number of non-adaptive, hitchhiker mutations, not all mutations can be implicated in resistance. However, if the same gene is mutated during adaptation to polymyxin in different studies that use different experimental conditions, it is more likely an adaptative allele than a hitchhiker. We identified such genes that were found to be mutated in our study as well as previous studies and arranged them in the following functional groups: two component system, *pmrAB*, lipopolysaccharide modification and biosynthesis genes (*migA, pagL* and *PA5194*), long chain fatty acid CoA-ligase (*fadD2*), outer membrane protein (*opr86*), probable short chain dehydrogenase *(PA4089)* and multidrug efflux transporter (*mexB*) While the role of some of these genes in polymyxin resistance has been validated (for example, *pmrAB, opr86, PA5194, pagL* and *mexB*), others have not been previously associated with resistance (12, 13, 27, 30, 33, 34). The targets were mapped on the phylogenetic trees (Fig. 3) to visualize the diversity and distribution of mutations among the end point isolates. The adaptive trajectories of these mutations are shown in Figures S3 and S4. Table 1 provides a list of candidates identified as putative players in colistin resistance among end point isolates and Fig. S5 shows their cellular localization.

### The number and variety of mutations observed in pmrB suggest that only modest changes in PmrB function are required for increased colistin resistance

PmrB is the sensor kinase of a two-component system, PmrAB and is involved in sensing cationic antimicrobial peptides (CAPs) (29). It was noteworthy that during the course of adaptation to colistin, 19 independent mutations were detected in *pmrB* using a 5% frequency cut-off for mutation detection. 18 of these mutations were SNPs that led to amino acid modifications affecting all the domains within this protein while one mutation was a 3 bp deletion leading to the loss of amino acid 47 in PmrB. Out of the 19 mutations, three (L167P, L170P and F408L) were observed independently in duplicate experiments and thus, 16 unique mutations were identified in *pmrB.*

Figure 5 shows the putative relationship of PmrB based on canonical sensor kinases of two-component systems. From the positions of mutations shown in red in Figure 5 (b), it is clear that every domain of PmrB was a potential target for adaptive mutations in this study. Also identified on this figure are adaptive mutations identified previously in clinical and lab-adapted *P. aeruginosa* strains (indicated in purple and black) highlighting the plasticity of the gene encoding this protein to acquire mutations. The propensity of *pmrB* to accumulate mutations at many locations suggests that modest changes in PmrB function are sufficient to alter the expression of genes in the regulon of this two-component system sensor kinase and confer resistance.

**Figure 5:**
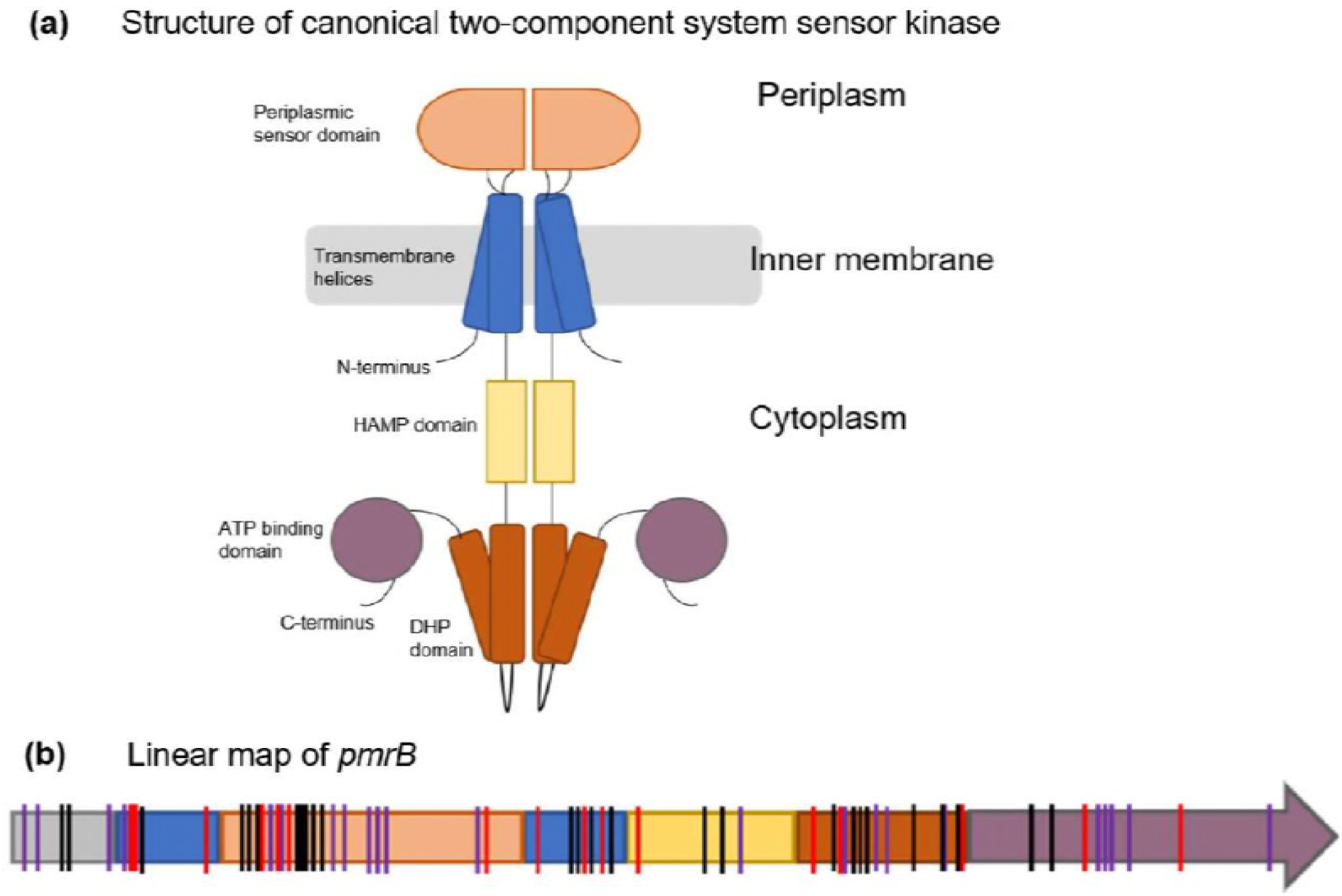
(a) Structural representation of a canonical two-component system sensor kinase dimer (adapted from (47)) and (b) linear map of *pmrB* showing positions of identified mutations. The color scheme for domains used in (a) has been maintained in (b). Mutations identified in *pmrB* in this study as well as previous works have been indicated by vertical lines on the *pmrB* gene (b). Red lines represent mutations identified in this study. Black lines represent mutations observed in other evolution experiments (12, 13, 27, 30) and purple lines represent mutations identified in colistin resistant clinical *P. aeruginosa* isolates (10–12, 26, 31, 32, 48). HAMP: histidine kinases, adenylate cyclases, methyltransferases and phosphodiesterases; DHP: dimerization and histidine phosphotransfer. Domain assignments in the PmrB protein are based on the predicted domain structure of PmrB of Moskowitz et al. (26). Details of the types of mutations observed in each domain of PmrB in this study can be found in Supplementary file S1.

### Introduction of *pmrB* mutations in a wild type PAO1 background indicates that other mutations are needed to explain the high levels of resistance

Multiple adaptive mutations identified in colistin resistant end point isolates suggested the possibility of clonal interference where multiple beneficial mutations in the population were competing with one another for success in the population (20). Since the role of PmrB in CAP resistance is known, we wanted to determine if specific changes in PmrB were sufficient to explain the very high MICs of some of the bioreactor end point isolates as well as clinical isolates from previous studies (26). Allelic replacement was used to identify the role of individual *pmrB* mutations in resistance by creating point mutations in *pmrB* within a wild type PAO1 background. 5 such constructs were made, each containing one *pmrB* mutation identified in the colistin evolved PAO1 populations from this work (L17P, L18P, D47G, L243R and F408L). These constructs and their respective bioreactor derived end point isolates were tested to compare colistin MICs (Table 3). Three *pmrB* mutations were observed in end point isolates that had been selected for whole genome sequencing – L18P, L243R and F408L. Sanger sequencing was used to identify the sequence of *pmrB* in a few other end point isolates and 2 isolates, labelled as #11 and #17 were selected that had the D47G and L17P mutations in PmrB, respectively. There is no information regarding other mutations in these isolates since whole genomes of these isolates were not sequenced.

**Table 3.**
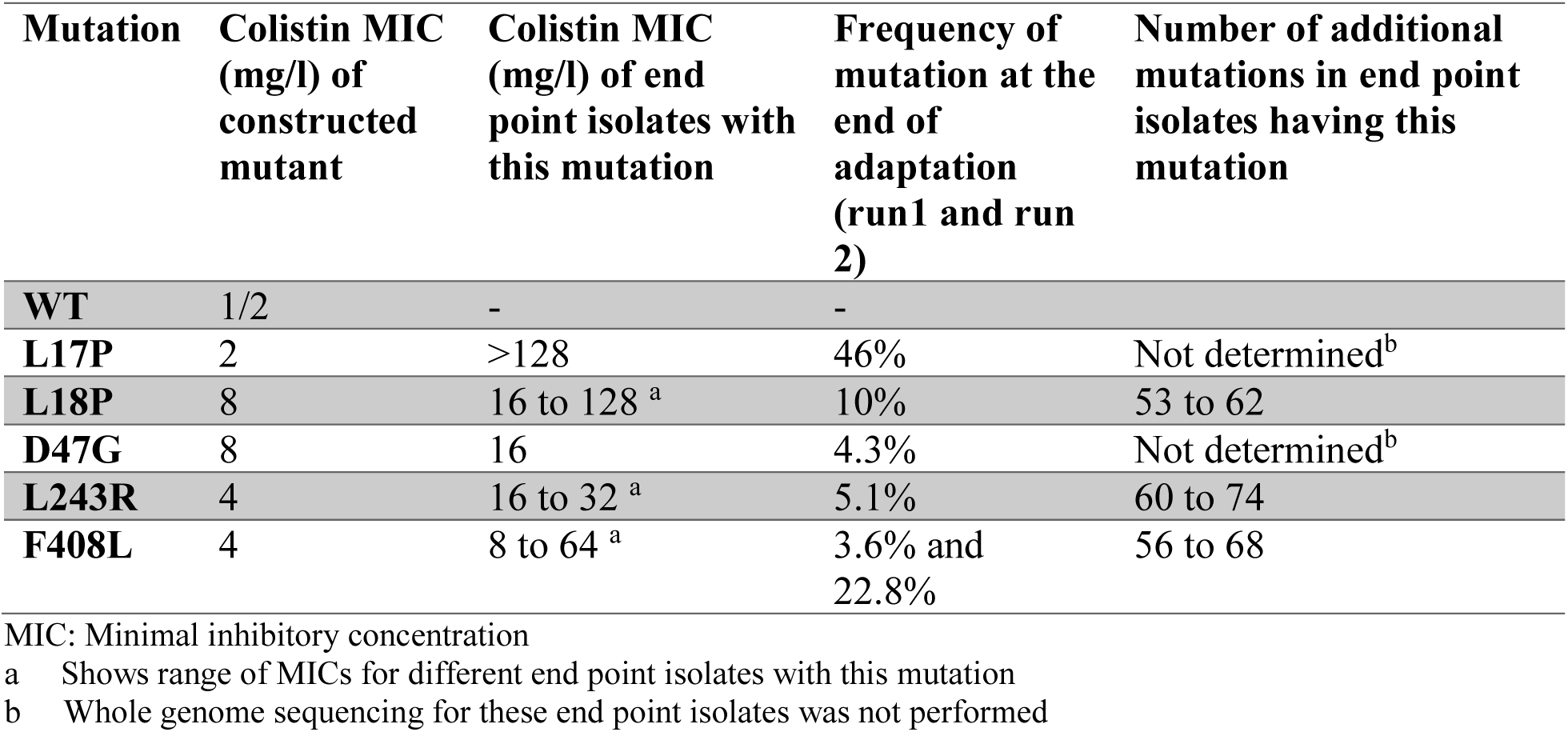
Comparison of constructed *pmrB* mutants and bioreactor derived end point isolates having the same *pmrB* mutation

| Mutation | Colistin MIC (mg/l) of constructed mutant | Colistin MIC (mg/l) of end point isolates with this mutation | Frequency of mutation at the end of adaptation (run1 and run 2) | Number of additional mutations in end point isolates having this mutation |
| --- | --- | --- | --- | --- |
| WT | 1/2 | - | - |  |
| L17P | 2 | >128 | 46% | Not determined<sup>b</sup> |
| L18P | 8 | 16 to 128 <sup>a</sup> | 10% | 53 to 62 |
| D47G | 8 | 16 | 4.3% | Not determined<sup>b</sup> |
| L243R | 4 | 16 to 32 <sup>a</sup> | 5.1% | 60 to 74 |
| F408L | 4 | 8 to 64 <sup>a</sup> | 3.6% and 22.8% | 56 to 68 |

MIC: Minimal inhibitory concentration
<sup>a</sup> Shows range of MICs for different end point isolates with this mutation
<sup>b</sup> Whole genome sequencing for these end point isolates was not performed

From Table 3, it is evident that different mutations within *pmrB* contribute to different levels of colistin resistance. While a mutation constructed in the transmembrane domain of PmrB, L18P imparted complete resistance to colistin (MIC 8 mg/l), adjacent mutation L17P did not (MIC 2 mg/l). The mutation L18P first appeared on day 18 of adaptation (Run 1) while L17P was seen only during the final day of adaptation when the population was growing at 16 mg/l colistin (Fig. S4). While early mutations during adaptation are often the most beneficial, additional mutations conferring typically smaller advantages can arise later (35). Appearance of L18P earlier during adaptation provided resistance to members of a population that was still evolving suggesting that it might be a primary mutation. L17P, which appeared later when the population was already growing at a high drug concentration (16 mg/l) was not an early primary mutation but may have played a role in enhancing the level of resistance in a specific genomic background that had achieved initial success (12). Other point mutations in the periplasmic domain (D47G), dimerization and phosphotransferase (L243R) and in the C-terminal ATP binding domains (F408L) all imparted resistance to colistin with the highest MIC at 8 mg/l while their corresponding bioreactor derived end point isolates consistently had acquired higher levels of resistance. This data provides strong evidence for the role of epistasis in high resistance of the bioreactor derived end point isolates.

### PAO1 incurs a fitness cost as a trade-off to acquiring colistin non-susceptibility

Growth characteristics of the constructed mutants and bioreactor end point isolates possessing the same mutation shed light on the fitness of the mutants in the presence and absence of colistin (Fig. 6). The growth of the ancestor, PAO1 served as the reference (Fig. 6 (a)). It was clear from our data that higher levels of colistin resistance, that were associated with hypermutation, led to reduced fitness in the isolates.

**Figure 6.**
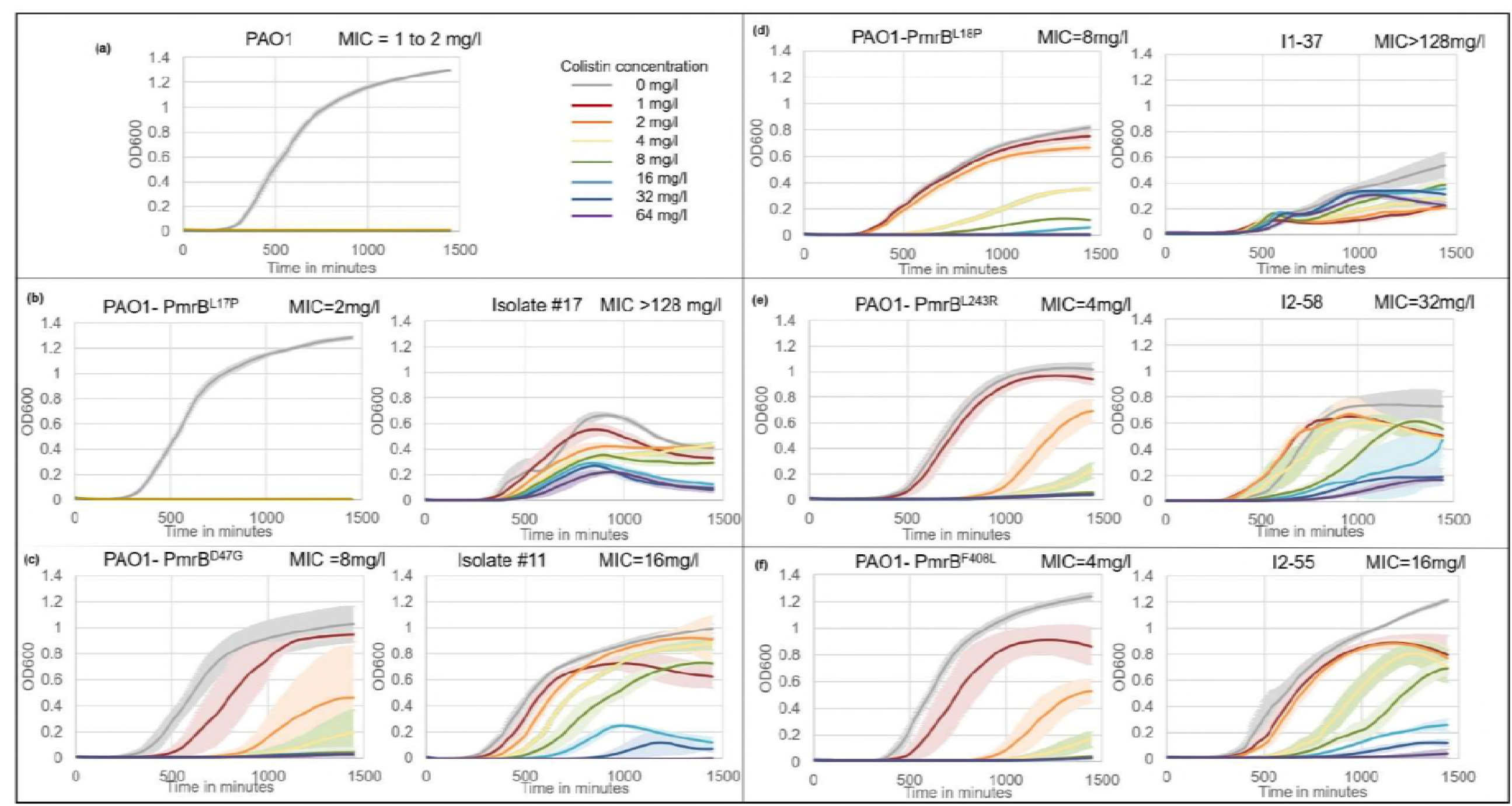
Growth characteristics of PAO1 ancestor, constructed point mutants and bioreactor adapted end point isolates. Each panel (except (a)) represents the constructed mutant on the left and the bioreactor adapted end point isolate carrying the same *pmrB* mutation on the right. All growth assays were conducted at colistin concentrations ranging from 0 to 64 mg/l. Error bars represent the standard deviation of three biological replicates. MICs of all these strains, measured by broth microdilution, are on the top right corner of each graph.

*pmrB* point mutants constructed in a wild type background could resist up to 8 mg/l colistin without undergoing a severe growth defect in the absence of the drug. In comparison, bioreactor isolates with the same *pmrB* mutations that had higher levels of resistance had decreased fitness in the absence of the drug. Bioreactor isolate I1-37 (doubling time = 397±90 minutes; MIC >128 mg/l) grew more than two times slower than the point mutant PmrB^L18P^ (doubling time = 165±5 minutes; MIC = 8 mg/l) in the absence of colistin (Fig. 6 (d)). Isolate I1-37, that had 59 mutations in addition to PmrB^L18P^ also had an increased lag time (longer by approximately 125 minutes) and lower overall yield but had the ability to survive in the presence of higher levels of colistin. Similarly, bioreactor isolate I2-55 (MIC = 16 mg/l) had a two-fold decrease in growth rate compared to the point mutant PmrB^F^408^L^ (MIC = 4 mg/l) (Fig. 6 (f)). However, it was capable of achieving nearly the same final cell density as PmrB^F^408^L^ as well as the ancestor strain (Fig. 6 (a)) suggesting that higher levels of resistance, as seen in I1-37, were associated with greater fitness defects. Although PmrB^L^17^P^ alone offered no adaptive advantage (Fig. 6 (a)), bioreactor derived isolate #17 that had a very high colistin MIC also had a severe growth defect.

The balance between acquisition of resistance and the associated fitness costs was further supported by allelic replacements to produce PmrB^D^47^G^ and PmrB^L^243^R^ in the wild type PAO1 background. These adaptive mutants were only modestly less fit in terms of growth rate and yield (Fig. 6 (c), (e) and (f)) under non-selective conditions but were able to grow better than wild type PAO1 in the presence of colistin which is also true for their corresponding bioreactor derived end point isolates.

Our data suggests that while *P. aeruginosa* acquires myriad mutations to resist colistin, the accumulation of these mutations comes at a fitness cost to the cells and higher levels of resistance are usually accompanied by a more severe growth phenotype in hypermutators. The advantage of mutation supply in a hypermutator is off-set by the fitness defect of the evolved isolates. In spite of the fitness cost, resistance to this drug of last resort in *P. aeruginosa* clinical isolates has been observed (36–38) which underscores the importance of understanding the mechanism of colistin resistance for the design of new strategies that can circumvent this problem.

## Discussion

The development of a hypermutator phenotype is a common occurrence in clinical settings and can lead to rapid adaptation to antibiotics (2, 14). Hypermutators increase the mutation supply within the evolving population allowing natural selection to act upon a more genetically diverse population thereby increasing the probability that a successful, e.g. a more antibiotic resistant variant can be found (18). The increased mutation load however comes with a cost to overall fitness as non-adaptive or hitchhiker mutations accumulate within the genomes of the hypermutators (15). As a random mutation is much more likely to be deleterious, the accumulation of these random mutations brings consequences to fitness especially when the selection pressure of the antibiotic is removed. For example, end-point isolates with increased MICs to colistin had substantially decreased growth rates in the absence of colistin (Fig. 6). While an increased mutation rate may be a poor long term evolutionary path for organisms like *P. aeruginosa*, the short-term benefit is clear. Interestingly, in our bioreactor environments that strongly favor the formation of biofilms, susceptible and non-hypermutator cells persisted despite the vessel containing >2 mg/l colistin for two weeks in the case of Run 1 (Fig. 2). We speculate that the biofilms insulate these weaker variants (Supplementary file S1) and can act as a reservoir for re-establishing a more fit population if the colistin were withdrawn. In an infected individual such as a cystic fibrosis patient, such a reservoir could suggest that when the antibiotic is switched, the more fit *P. aeruginosa* strains can re-emerge and conversely that colistin resistant variants could now be a reservoir for re-establishing the colistin resistant population if the patient returned to colistin therapy. Thus, the combination of hypermutation and strong biofilms can lead to the persistent and difficult to treat infections that are a hallmark of *P. aeruginosa.*

It has been suggested that because of the large number of hitchhiker mutations that succeed in the population under conditions of selection, the signature of selection in the genome is very weak, making it difficult to distinguish driver alleles from passenger mutations (15). We show that it is possible to identify the signature of selection in an adapting hypermutator population using a combination of genomic and statistical approaches. Quantitative experimental evolution provides a ready means to construct the genomic data needed for analysis (13, 16, 19–21). We propose a hierarchy of analyses beginning with the Fisher’s Exact Test of both end-point isolates and longitudinal metagenomic data to build and rank a list of candidate genes that are putatively involved in resistance. This method has proven useful in identification of adaptive mutations in previous studies involving hypermutation and weak selection (16, 19). We also construct phylogenetic trees of the end-point isolates that provide further genetic and evolutionary structure to the candidate list. Furthermore, we use the information about the frequencies of these mutations in the daily populations to build parsimonious evolutionary trajectories for the most important drivers for colistin resistance and taken together these trajectories illustrate what we term the “adaptive genome” of *P. aeruginosa* to the selection environment (Fig. 7).

**Figure 7.**
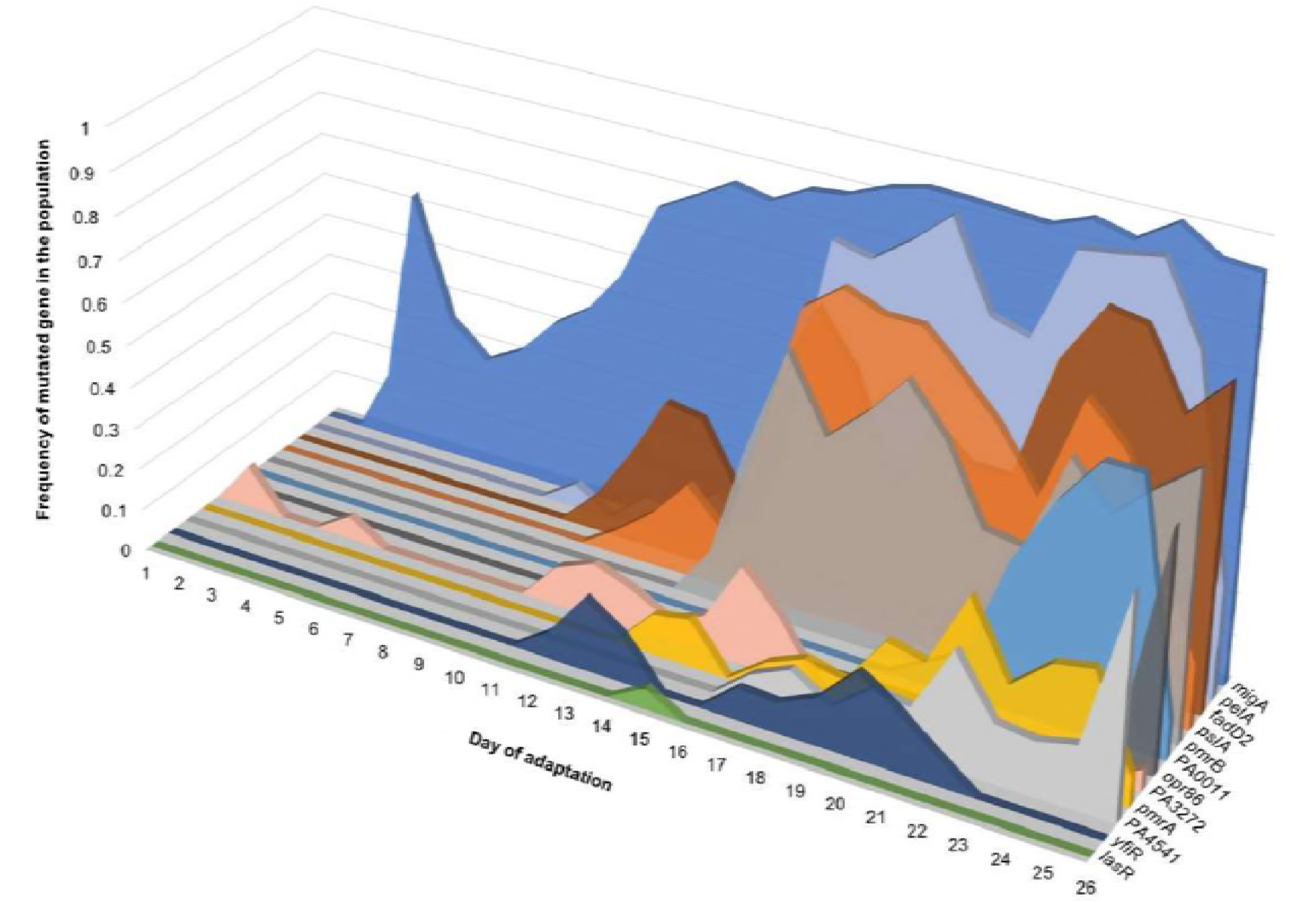
Adaptive genome of a *P. aeruginosa* population (Run 1) evolving to colistin. Most adaptive genes had multiple mutations within them during the course of adaptation. On a given day, the sum of frequencies of the different alleles within a gene was calculated and plotted on the Y-axis. The X-axis represents the day of adaptation and the Z-axis represents each gene. Genes represented here include putative candidates listed in Table 1 and selected candidates from Table 2 (selection based on their appearance in other polymyxin resistance studies).

Using genetics and phenotypic analysis of growth rates, we establish that epistatic interactions between multiple mutations in the bioreactor derived hypermutator end point isolates are most likely responsible for the observed higher levels of colistin resistance which comes at a fitness cost to the cells in terms of reduced growth rate and overall yield (Fig. 6). Such epistatic interactions between multiple alleles implies that high levels of colistin resistance can be acquired via multiple adaptive routes which in turn result in the formation of a rugged fitness landscape with many local peaks, each representing a local optimum. Hypermutation provides an effective means for the cells to access this landscape during selection.

In summary, this work sheds light on multiple features of *P. aeruginosa’s* evolvability to the drug of last resort, colistin. While the complexities of hypermutation have hindered progress, it has become increasingly clear that modern next-gen sequencing and experimental evolution provide a path forward to the study of this clinically and conceptually important evolutionary mechanism for adaptation (13, 16, 39). The methodological approaches within this work show that hypermutation can be studied successfully to produce a broad understanding of adaptive evolution in established as well as in future emerging pathogens.

## Materials and methods

### Bacterial strains, plasmids and growth conditions

*P. aeruginosa* PAO1 was obtained from American Type Culture Collection (ATCC 15692). Plasmid pEX18Gm was kindly provided by Dr. Herbert Schweizer. PAO1 was routinely grown in Lysogeny Broth (LB: 10 g/l tryptone, 5 g/l yeast extract, 10 g/l sodium chloride) or on LB + 15 g/l bacto agar. Growth medium for adaptation of PAO1 to colistin was LBHI (80% LB + 20% brain heart infusion (BHI) medium) supplemented with 2 mM magnesium sulfate and appropriate concentration of colistin. Colistin stock solution was made by dissolving colistin sulfate (DOT Scientific Inc., MI, USA) in water followed by filter sterilization using a 0.22 µm filter. Cation adjusted Mueller Hinton broth (CA-MHB) was used for minimum inhibitory concentration (MIC) testing and growth curves. Gentamicin at 20 mg/l was used for maintenance of pEX18Gm in *Escherichia coli* and 60 mg/l was used for growing PAO1 transformed with the pEX18-*pmrB* plasmids. Strains and plasmids used in this study are listed in Table 4.

**Table 4.**
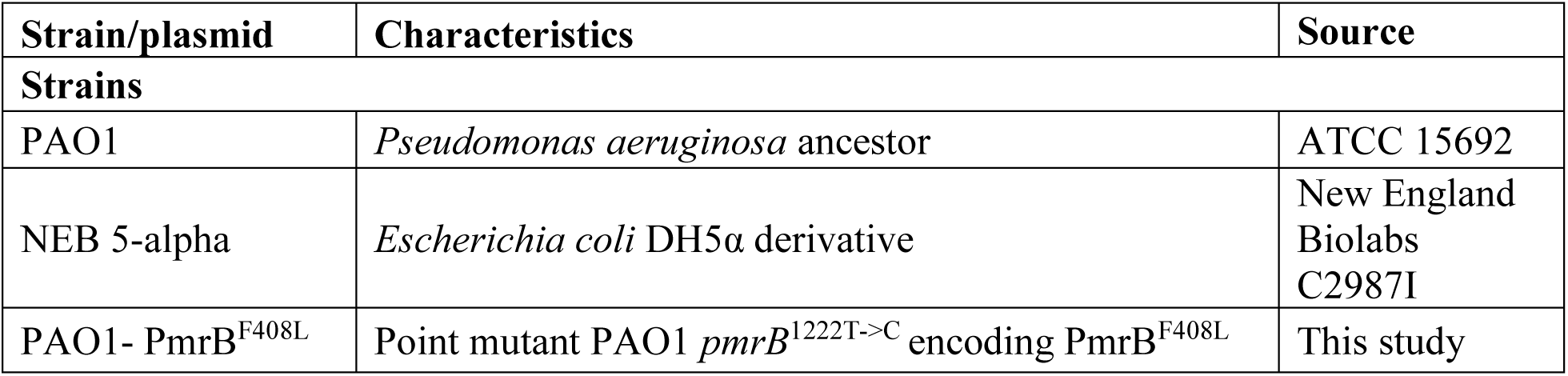
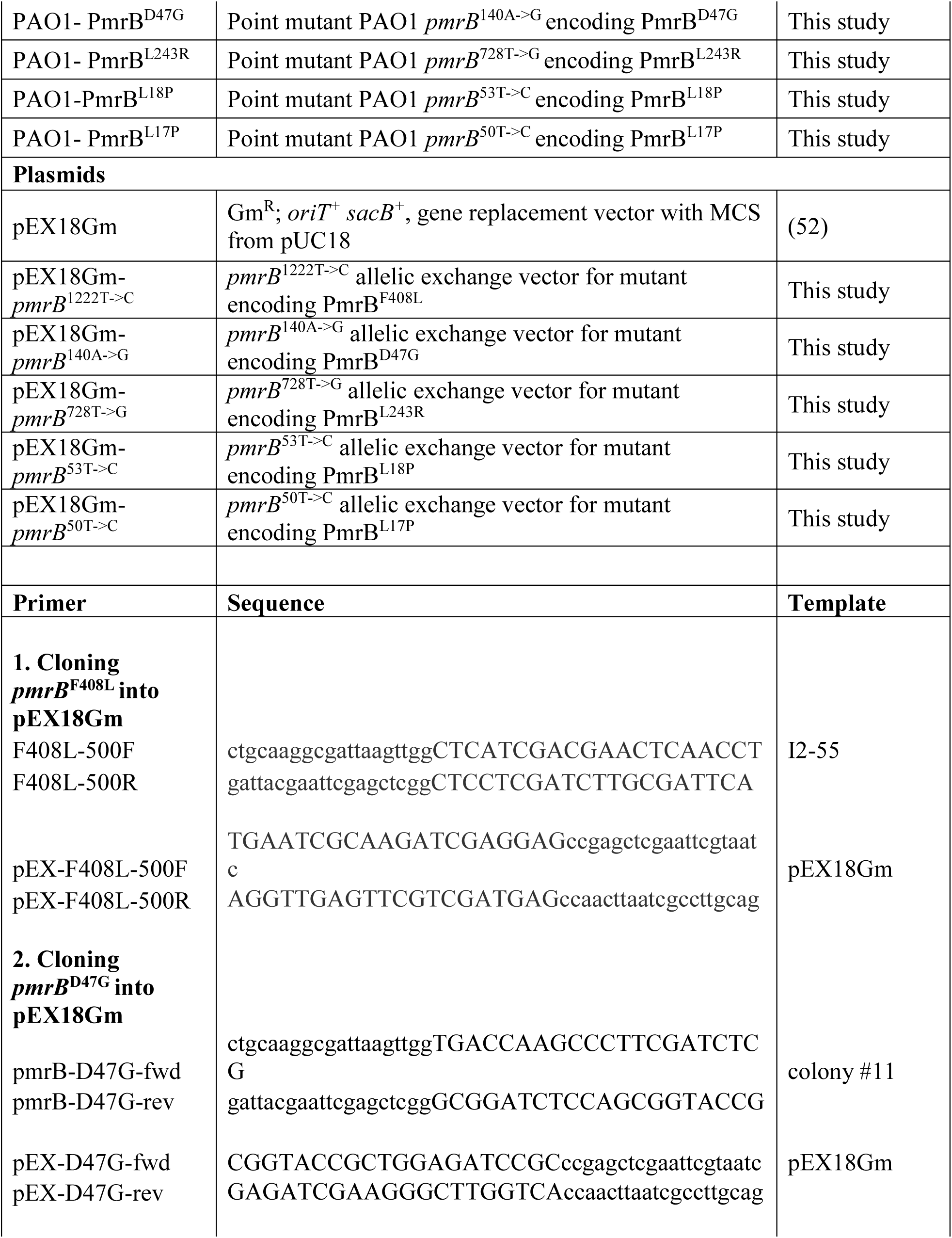
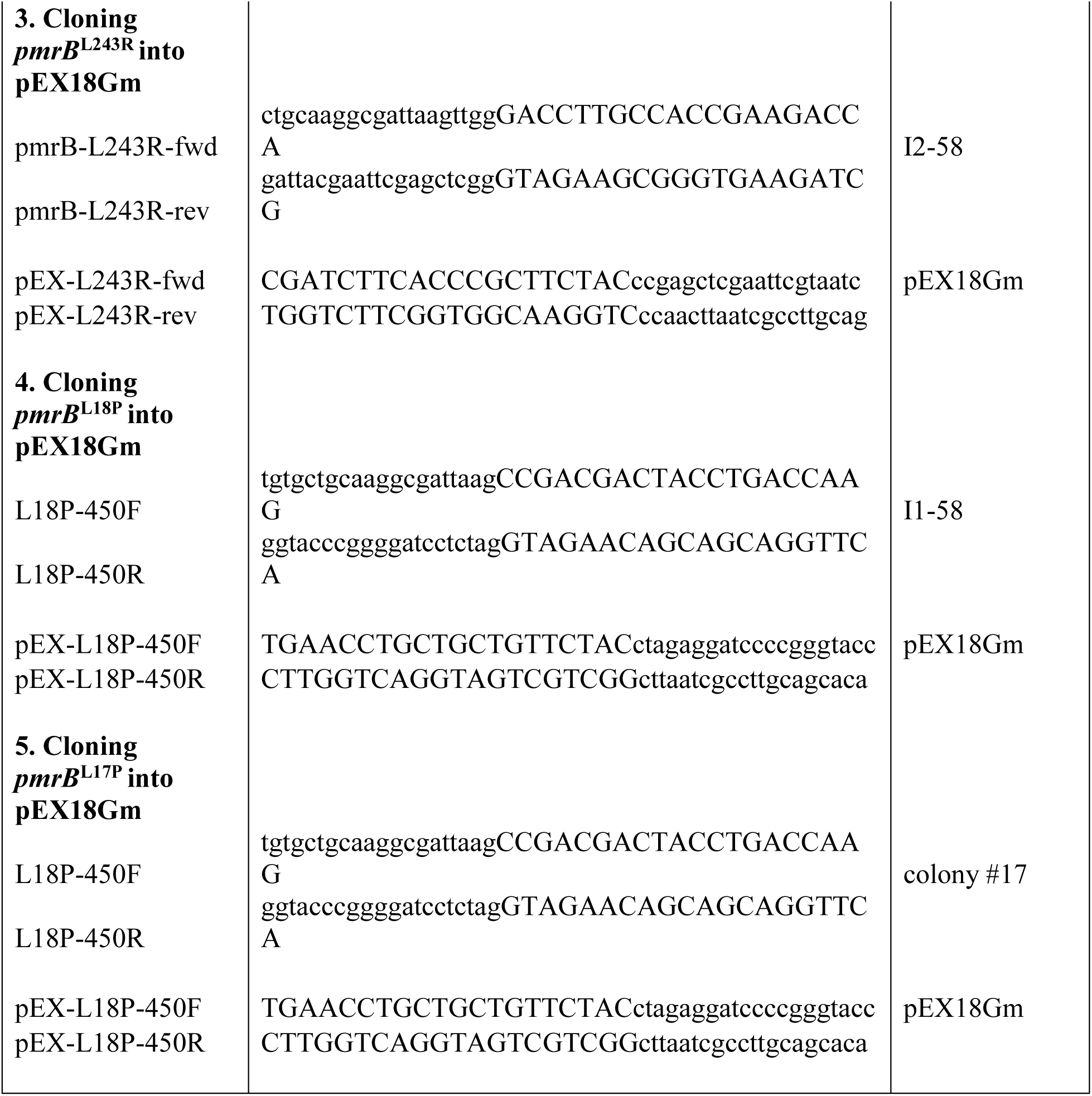
Bacterial strains, plasmids and primers used in this study.

### Evolution of PAO1 to colistin

PAO1 was evolved to colistin in a modified turbidostat as described in (24). In duplicate runs, a 300 ml PAO1 culture was established in the bioreactor vessel by using a single colony as inoculum. The culture was maintained in mid-exponential growth phase using respiratory CO_2_ as a proxy for turbidity to control media flow. After 12 hours of growth, the first sub-inhibitory dose of colistin was added to the vessel (0.5 mg/l). After that, the culture was monitored and the drug concentration was empirically increased. Details of the process are provided in (24). PAO1 was able to grow at 18 mg/l colistin after 26 days of evolution during run 1 and at 16 mg/l colistin after 17 days of evolution during run 2.

### Isolation and characterization of end point isolates

The final resistant population of PAO1 was serially diluted and spread on non-selective medium (LBHI + 2mM magnesium sulfate) to isolate individual members of the population. Each colony was called an end point isolate. 88 end point isolates were selected from run 1 and 82 from run 2 for further phenotypic characterization. Morphological characteristics were recorded for each isolate which included colony size, shape, color, consistency and appearance when grown in a non-selective liquid medium (Supplementary file S1). Minimum inhibitory concentration (MIC) of colistin was tested using a preliminary broth microdilution assay. A colistin gradient (0 to 128 mg/l) was set up in a 96 well polypropylene plate containing 100 µl CA-MHB per well. 1 µl of an overnight culture (grown in LBHI + 2mM magnesium sulfate) for each isolate was used as inoculum. Plates were incubated at 37°C and visible growth was recorded after 20-24 hours. The lowest colistin concentration showing absence of growth was the MIC. MIC of other antibiotics were also tested for determining cross-sensitivity/cross-resistance to other drugs. Agar dilution MIC assays were performed for this in 100 mm petri dishes containing CA-MH agar with appropriate drug concentration. A 96 pin applicator was used to spot the overnight culture of each isolate on the agar plates. MICs were recorded after 20-24 hours of incubation at 37°C. Based on the different phenotypic traits, 15 end point isolates from run 1 and 14 from run 2 were selected for whole genome sequencing and mutation identification. Cross sensitivities to other antibiotics were observed in some end point isolates but a correlation of the cross sensitivity to colistin resistance could not be established and hence, this phenotype was not pursued further.

### Minimum inhibitory concentration assay

Broth microdilution MIC tests in biological triplicate were performed for the 29 end point isolates selected for whole genome sequencing using 96 well polypropylene plates. Each well was filled with 100 µl CA-MHB and appropriate concentration of colistin (0-128 mg/l colistin in 2-fold increments). Overnight cultures for end point isolates were grown as biological triplicate in LBHI + 2 mM magnesium sulfate. Optical densities of the cultures were adjusted to 0.05 and 5 µl of this OD adjusted culture was used to inoculate each well of the plates. Growth was checked visually after 20-24 hours of incubation at 37°C and MICs were recorded.

### Whole genome sequencing and analysis

DNA isolation and whole genome sequencing was performed as described in (24). Samples from run 1 of adaptation were sequenced at the US Army Edgewood Chemical Biological Center (ECBC, MD, USA) as 100 bp paired end reads and samples from run 2 were sequenced by a commercial facility (Genewiz, NJ, USA) as 150 bp paired end reads. Read trimming of raw Fastq reads was performed using Sickle (40). Trimmed reads were analyzed using Breseq version 0.30.1 (41) to identify genetic variations between ancestor and adaptive populations as well as end point isolates. The genome sequence of the ancestor was obtained from NCBI (AE004091). The ancestor colony used to inoculate the bioreactor during each run was re-sequenced and the APPLY function on Breseq was used to incorporate any differences in the genome of the re-sequenced ancestor strain into the NCBI reference genome before using it for identifying mutations in the evolved strains. The consensus mode was used for identification of mutations in the end point isolates. The polymorphism mode on Breseq was used for the analysis of daily metagenomic populations from the bioreactor using the following command: -p -- polymorphism-reject-surrounding-homopolymer-length 5 --polymorphism-reject-indel-homopolymer-length 0 --polymorphism-minimum-coverage-each-strand 6 --polymorphism-frequency-cutoff 0.02. To filter low quality mutations from the daily populations under analysis, two additional quality-filtering steps were added after the Breseq analysis of each daily sample. It was observed that the number of false calls increased substantially for low frequency mutations. Thus, mutations in the daily populations that fell below a threshold of 5% frequency in the population were filtered out. Second, it was found that mutations had been called in several reads that had low Mapping Quality (MQ) scores. Mutations in regions that contained three or more reads having an MQ score less than 100 were also filtered out. Also, reads were manually examined and mutation calls which were characterized by 3 or more mutations clustered within a read at low frequencies and whose occurrence was not consistent with adaptation or hitchhiking were eliminated from further analysis. Dataset S1 contains a list of all mutations and their frequencies on each day of evolution during Runs 1 and 2.

### Construction of phylogenetic trees

The Breseq output for each end point isolate contained a list of mutations in the genome of the end point isolate. The APPLY function on Breseq was used to apply the mutations in the end point isolate onto the PAO1 reference genome to create a genome sequence for each end point isolate. Next, the genome of the reference strain was manually aligned with that of all the end point isolates using the software MEGA7 (42). Phylogenetic trees were constructed for both runs using the maximum parsimony algorithm based on (43) as implemented in MEGA7. These trees were then visualized using the Dendroscope3 software (44).

### Fisher’s Exact Test

To perform the Fisher’s Exact Test on end point isolates, a list of the total number of mutations occurring in a gene was compiled from the Breseq output file. Gene lengths were obtained from the Pseudomonas Genome Database (45) by mapping the gene name to the gene length. A few genes without matches were manually given gene lengths. With the compiled list containing the number of mutations per gene, the length of the gene, the total number of mutations in all end point isolates or populations and the total length of the PAO1 genome, a Fisher’s Exact Test was performed using the fisher.test function in R with a “two sided” alternative hypothesis and a significance threshold of 0.001.

### Construction of point mutations in PAO1 *pmrB*

Point mutants in PAO1 were constructed using the protocol described in (46) with minor modifications. pEX18Gm was used instead of pEX18Tc. Allelic exchange vectors (Table 4) were made using Gibson Assembly^®^ Master Mix (New England BioLabs) with primers listed in Table 4. Mutant *pmrB* alleles were amplified from the respective bioreactor end point isolate containing the mutation. Electroporation of the constructed plasmid into PAO1 was done at room temperature using a 2mm gap electroporation cuvette at 2.2 kV as described previously (46). Electroporated cells were spread on BHI + 60 mg/l gentamicin (Gm) plates and incubated at 37°C for 2-3 days. Gm resistant colonies were colony purified on BHI + Gm60 plates and single colonies were streaked on no salt LB (NSLB) + 15% sucrose plates and incubated at 30°C for sucrose counterselection. Sucrose resistant colonies were tested for Gm sensitivity and for colonies that were Gm sensitive and sucrose resistant, the *pmrB* gene was amplified and sequenced using Sanger sequencing to determine presence of the point mutation.

### Growth curves

Overnight cultures were prepared in LB broth in biological triplicate. Optical densities were measured and normalized to 0.05. 96 well polypropylene plates were used for growth curves. Each well was filled with 100 µl CA-MHB and a colistin gradient was set up for concentrations ranging from 0 to 64 mg/l. Each well was inoculated with 1 µl of the OD normalized culture and growth was measured in each well using a BioTek Epoch2 microplate reader at 37°C for 24 hours with optical density being measured at 5-minute intervals. The OD of plain CA-MHB (blank reading) was subtracted from the OD of the samples at every time point. Exponential smoothing was applied to each data series to account for the noise in the OD measurements due to clumping and biofilm formation in the wells. The final graph of OD versus time was plotted using the average of 3 biological replicates with standard deviation. Doubling times were calculated in the OD_600_ interval 0.2 - 0.4.

## Acknowledgements

We are thankful to Dr. Luay Nakhleh for his assistance with computational analysis and resources. We are thankful to Dr. Herbert Schweizer for kindly providing plasmids for genetically modifying *P. aeruginosa.* We appreciate the advice provided by Weiliang Huang and Dr. Angela Wilks for isolation and identification of mutant *P. aeruginosa* candidates following allelic replacement. We are grateful to Dr. Jeff Barrick for assistance with analysis of metagenomic population data using Breseq.

## Data availability

Whole genome sequencing data generated during this study are submitted to the Sequence Read Archive (SRA) database under the BioProject Accession number PRJNA486960.

## Author contributions

. S. conceived of the idea. H. H. M., A. P. and K. B. conducted the bioreactor adaptation experiments. M. K. and H. S. G. conducted whole genome sequencing for Run 1 of this project. H. H. M. was involved in analyzing the WGS data and in mutation identification. R. A. L. E. performed the phylogenetic and statistical analysis. H. H. M. performed the phenotypic characterization and allelic replacements. H. H. M. and Y. S. wrote the manuscript. All the authors provided critical feedback and contributed to the manuscript.

## Materials and correspondence

Material requests and correspondence should be addressed to Professor Yousif Shamoo.

## Supplemental material legends

**Figure S1**. Phenotypic diversity of end point isolates obtained at the end of adaptation of PAO1 to colistin. (a) End point isolates showed variations in the size, color and texture. (b) Pie chart showing colistin minimum inhibitory concentrations (MICs) of end point isolates. Numbers in each segment represent the actual number of isolates from that population having the specific colistin MIC as indicated by the colors in the legend. 12 out of 88 isolates from run 1 (14%) and 19 out of the 82 isolates from run 2 (23%) were colistin susceptible (MIC ≤ 2 mg/l).

**Figure S2**. (a) Excision and circularization of phage Pf4 upon colistin exposure. The prophage, Pf4 exists in lysogenic state in PAO1. During exposure to colistin, the phage encoded DNA excised from the PAO1 chromosome and formed superinfective phage. (b) Induction of prophage during evolution of PAO1 to colistin. Top left panel shows different dilutions of the supernatant from day 1 of evolution (before drug exposure) that are incapable of lysing the lawn of PAO1 on the plate. The supernatant obtained after centrifugation of the population sample was filter sterilized and then serially diluted (10-fold dilutions) in SM buffer (50 mM Tris-HCl, pH 7.5 + 100 mM NaCl + 10 mM MgSO_4_.7H_2_O). 5 µl of each dilution was spotted on a lawn of PAO1. Supernatant from day 2 (first instance of drug exposure) has strong lytic activity (top right) suggesting induction of prophage and lytic capability. This lytic capability continues to exist till the end of adaptation (bottom panels -days 13 and 25). All these samples are from run 1 of adaptation that lasted 26 days.

**Figure S3**. Evolutionary trajectories of adaptive mutations identified in this study Each graph is a plot of the frequency of a mutation in a gene within the population versus the day of adaptation on which it was observed. Multiple mutations within a gene are plotted on the same graph using different colors to represent each mutation. Only alleles that rose above 10% frequency during adaptation are shown here. Evolutionary trajectories of the adaptive alleles of *pmrB* are shown in Figure S4.

**Figure S4**. Trajectories of *pmrB* mutations detected at ≥ 5% frequency during adaptation to colistin. 11 mutations were identified in run 1 which lasted 26 days. Out of the 11, only 3 mutations, L17P, L18P and L167P were detected in the final resistant population at ≥ 5% frequency. Run 2 which lasted 17 days had 8 *pmrB* mutations in the evolving population with only 2 mutations, L243R and F408L detectable in the final resistant population.

**Figure S5**. Cellular localization of targets identified in this study playing putative roles in colistin resistance. Targets in purple text were identified by the Fisher’s Exact test of end point isolates and targets underlined were common among our study and other polymyxin resistance studies.

**Figure S6**. Relationship between colistin MIC of an end point isolate and its biofilm forming capability as measured by crystal violet staining. No significant co-relation between level of resistance of the isolate and its biofilm forming capability can be inferred from this data.

**Table S1.** List of end point isolates selected for whole genome sequencing and their minimum inhibitory concentrations (MICs) to colistin

**Table S2**. Hypothetical protein encoding genes identified as significant by Fisher’s Exact Test performed on mutations in daily populations of PAO1 evolving to colistin

**Dataset S1.** Mutations identified in daily populations and well as in end point isolates obtained from adaptation of PAO1 to colistin during Run 1 and Run 2.

